# Genetic susceptibility to obesity-related asthma and its modulation by sequelae of obesity

**DOI:** 10.64898/2026.06.16.731769

**Authors:** David A. Thompson, Yvonne B. Wabara, Sarai Duran, Anna Reichenbach, Deepa Rastogi

## Abstract

**Rationale:** Asthma is a multifactorial disease with a role of genetic susceptibility and environmental exposures. These aspects are poorly understood for pediatric obesity-related asthma, a phenotype of non-allergic asthma.

**Objective:** To quantify gene by environment interactions in obesity-related asthma

**Methods:** Using expression quantitative trait loci (eQTLs) as measure of genetic susceptibility, and obesity-mediated effects on anthropometrics, metabolic measures, and T helper cell proportions as biological sequelae of obesogenic environment, we quantified the association of eQTLs with asthma burden, and its modulation by obesity-mediated effects, in primary cohort of 144 children, and validation cohort of 101 children.

**Measurements and Main Results:** Of the 3,904 eQTLs associated with gene expression, up to 30% were associated with pulmonary function indices, including FVC, FEV_1_, TLC, FRC and IC, and were enriched for African ancestry. These eQTLs encoded for antigen presentation, cell mobility, autophagy, small GTPase signal transduction, fatty acid metabolism, and chromosome segregation pathways. Neck and waist circumference, insulin resistance, leptin and adiponectin levels, and T helper 1 and 17 cell proportions attenuated association of up to 51% eQTLs with pulmonary function, which encoded for all but fatty acid metabolism and chromosome segregation pathways. eQTLs associated with *ATF6* and *MEI1* retained significance. eQTLs for *RNASET2, FBLN5, STX2, HEATR3 and SERPINB6* genes were associated with pulmonary function in the validation cohort.

**Conclusions:** We report novel genetic susceptibility markers of asthma burden in pediatric obesity-related asthma that are enriched for African ancestry and are partly attenuated by truncal fat load and obesity-mediated inflammation and metabolic dysregulation.

## Introduction

Pediatric asthma is broadly grouped into Type 2 (T2)-hi and T2-lo asthma, based on presence or absence of atopic or T helper (Th) cell type 2 mediated inflammation (1, 2). While T2-hi asthma is extensively investigated and has targeted therapies (3), T2-lo asthma, limited by the absence rather than presence of defining features, is less well understood. Among its diverse etiologies (4), obesity-related asthma is the better understood T2-lo phenotype that is the leading cause of incident asthma due to global increase in childhood obesity (5). Its association with higher disease burden and poor medication responsiveness (6) highlight the need to elucidate its pathobiology for targeted therapeutic development.

Asthma is a multifactorial disease with a role of genetic susceptibility and of environmental exposures (7). While T2-hi asthma is associated with single nucleotide polymorphisms (SNPs) in the *ORMDL3* gene that predispose to atopic inflammation and altered immune response to viral illnesses, a key environmental exposure associated with disease burden, similar details are lacking for T2-lo asthma such as obesity-related asthma. There is increasing recognition of genetic susceptibility to obesity-related asthma, including that related to ancestry (8–11). Obesity-mediated complications including truncal adiposity, metabolic abnormalities, and immune perturbations, that are biological sequelae of an obesogenic environment (12) also contribute to the burden of obesity-related asthma (13–15). However, the interplay between genetic susceptibility, including ancestry, and obesity-mediated complications in pathobiology and disease burden of obesity-related asthma is not known.

To address this knowledge gap, we quantified the association of genetic susceptibility, defined by expression quantitative trait loci (eQTLs), with disease burden of obesity-related asthma, given their functional relevance to multifactorial diseases via modulation of gene expression (16, 17), and the modifying effects of anthropometrics, metabolic measures, and Th cell proportions, on this association of eQTLs with obese asthma burden.

## Methods

Additional methodological details are included in the online supplement.

### Study population

The primary cohort was comprised of 144 children, ages 7-18 years, including 57 with obesity-related asthma (OA), 33 with healthy-weight-asthma (HwA), 27 with obesity-alone (Ob), and 27 healthy-weight without asthma (healthy-weight controls (HC)). The validation cohort comprised of 101 children, ages 7-11 years, included 48 with OA and 53 with HwA (18). Obesity was defined as body mass index (BMI) >95^th^ percentile for age and sex (19). Asthma was defined by physician diagnosis, active prescription of medications, and reversible airflow obstruction (20). The study was approved by the Institutional Review Board and informed consent was obtained at the study visit (21).

### Measures of asthma burden

Asthma burden was measured with pulmonary function indices and symptom-based classification of severity using Composite Asthma Severity Index (CASI) (22), and control using Asthma Control Test (ACT) (23, 24). Percent predicted values of FVC, FEV_1_, and FEV_1_/FVC ratio were calculated using race-neutral Global Lung function Initiative (GLI)(25), and percent predicted values of FEF_25-75%_, TLC, RV, RV/ TLC ratio, ERV, FRC, and IC were calculated using ATS/ERS guidelines (26).

### Demographic, anthropometric, and metabolic variables

Demographics included age, sex, race/ethnicity, and anthropometrics included height, weight, BMI percentile, and neck, midarm, waist, and hip circumference. Metabolic measures included fasting levels of glucose, insulin, lipids, and adipokines, leptin and adiponectin (13). Insulin resistance was defined by Homeostatic Measurement of Insulin Resistance (HOMA-IR).

### Th cell subsets

Blood CD4+T (Th) cells underwent flow cytometric analysis of cell surface staining with CD4, CD25, CD127, CXCR3, and CCR6 antibodies (Biolegend, San Diego, US) to quantify T helper cell subsets (Th1, Th1/17, naïve/Th2, Th17, and regulatory T cells) **[Figure E1]**.

### Transcriptomic analysis of CD4+ T cells

500 ng RNA from CD4+T cells underwent library preparation with KAPA stranded mRNA-Seq kit (KAPA Biosystems, Wilmington, MA) that were sequenced on NovaSeq 6000, and analyzed on DESeq2 to elucidate between-group differences in gene expression (21).

### Genotyping and eQTL mapping

CD4+T cell DNA was genotyped on the Infinium Multi-Ethnic Global Bead Chip (27). Cis eQTLs were variants within 1 Mb of a gene’s transcription start site associated with its expression. Analyses were limited to autosomal variants, removing those with a minor allele frequency (MAF) of less than 0.1, or those that did not follow the Hardy-Weinberg equilibrium (p-value<0.000001). Linkage disequilibrium (LD) within the final eQTL set was identified using SNP2GENE module from Functional Mapping and Annotation of Genome-Wide Association Studies (FUMA GWAS) (28).

### Statistical analysis

Analysis was conducted on R statistical software v. 4.4.1. We quantified the univariate association of eQTLs with asthma burden, and elucidated their biological relevance with Gene Ontology (GO) pathway analysis (29). Univariate association of anthropometrics, metabolic measures, and Th cell proportions with asthma burden was quantified using simple linear regression analysis. Effect of anthropometrics, metabolic measures, and Th cell proportions on association of eQTLs with asthma burden was quantified by multivariate linear regression analysis. This was followed by Similarity Network Fusion (SNF) multi-omics analysis (30) to cluster samples with high disease burden irrespective of study group. Ancestry was classified with ADMIXTURE v1.3.0 (31). eQTLs overlapping between primary and validation cohorts were investigated for their association with asthma burden in the validation cohort. Benjamini Hochberg method was applied to calculate the false discovery rate (FDR) (32).

## Results

### Cohort characteristics

Demographics did not differ between the study groups. Compared to HwA and HC groups, weight, BMI percentile, and neck, midarm, waist, and hip circumference, as well as triglycerides, insulin, HOMA-IR, and leptin were higher, and HDL and adiponectin levels were lower among OA and Ob groups. Naïve/Th2 were higher among OA compared to the other groups **[Table 1]**. Among measures of asthma burden, FVC and FEV_1_ did not differ but FEV_1_/FVC ratio was the lowest in OA relative to other groups, while RV/TLC ratio was lowest and IC was highest in Ob group. CASI and ACT did not differ between OA and HwA **[Table 1]**.

**Table 1.**
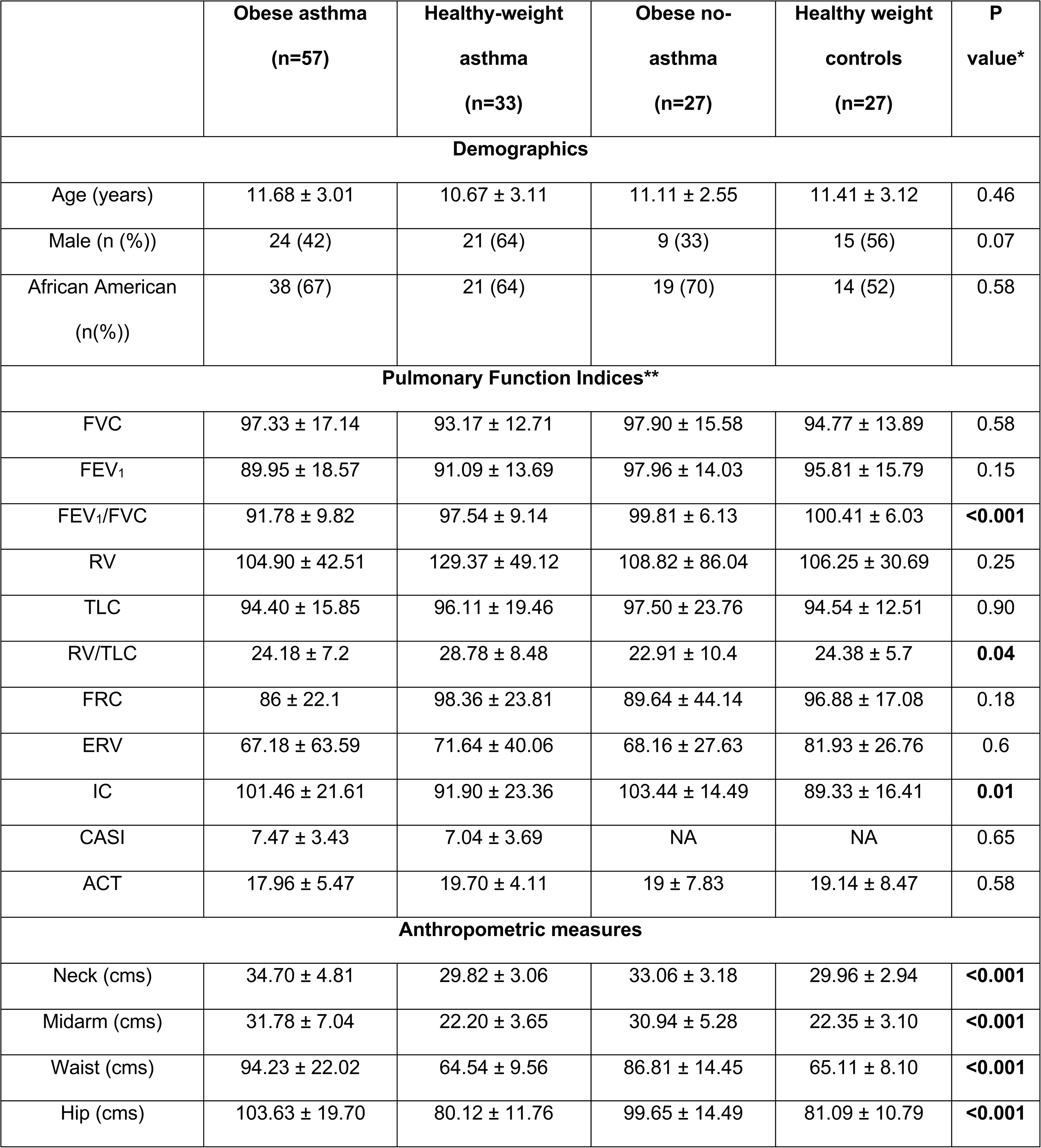
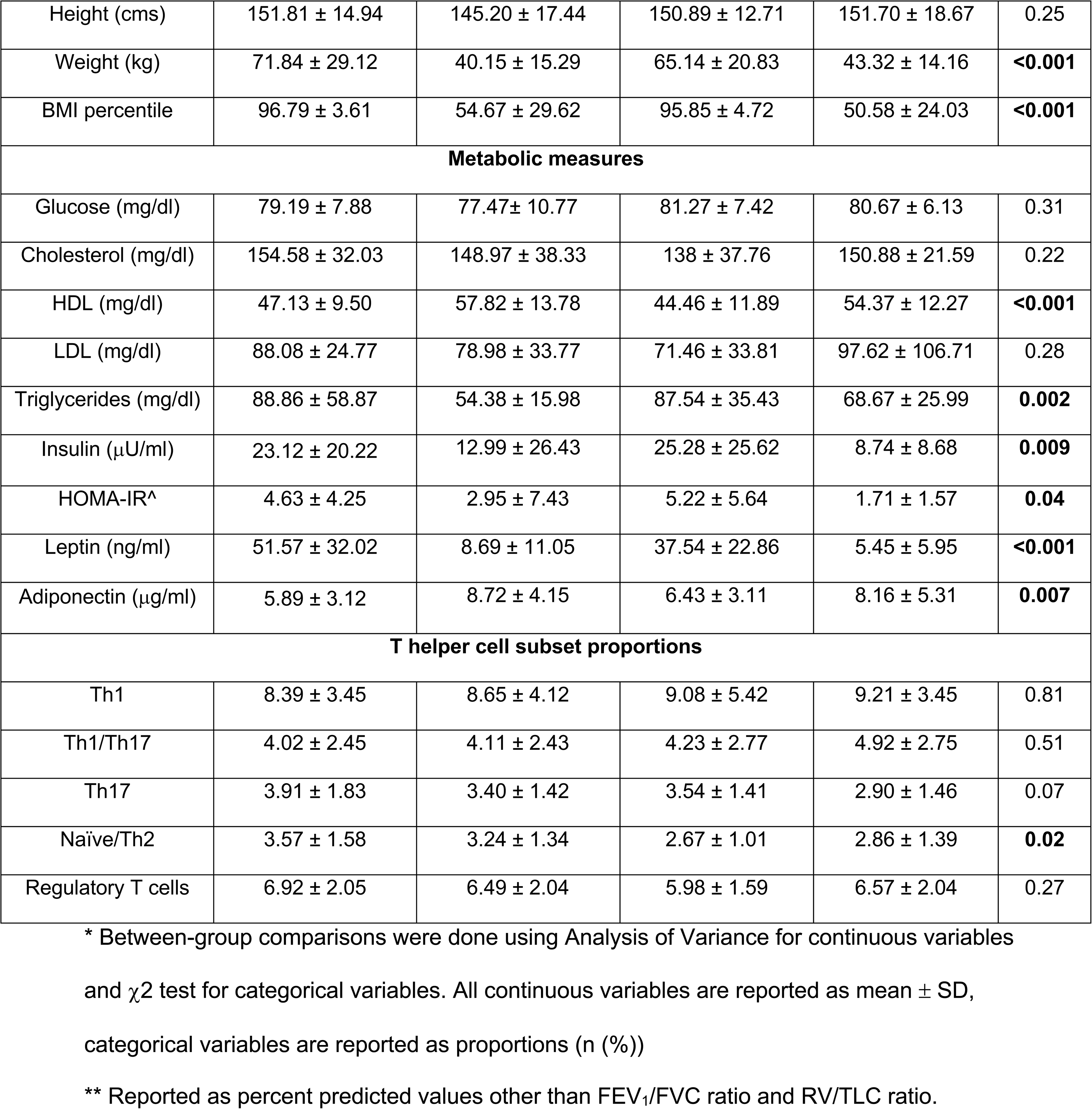
Demographic and clinical characteristics of study participants in primary cohort.

### Association of eQTLs with asthma burden

There were 8,284 eQTLs **[Table E1a]**, of which 75 were homozygous for major allele in all participants and excluded from further analysis. In the remaining 8,209 eQTLs **[Table E1b]**, FUMA analysis identified 13 lead SNPS in LD with 307 other SNPs, none of which were present in the 8,209 eQTLs. The MAF differed by race/ethnicity in 10.7% (n=878) eQTLs but did not differ by sex **[Table E1c]**. Among the 8,209 eQTLs, 18% (n=1,544) were associated with FVC, 29.4% (n=2,413) with FEV_1_, 15.2% (n=1,246) with TLC, 0.003% (n=22) with RV, 0.05% (n=408) with FRC, and 0.02% (n=124) with IC **[Table E2a-f]**, while none were associated with ERV, FEV_1_/FVC or RV/TLC ratio, CASI, or ACT. Limiting the analysis to 3,904 eQTLs associated with differential gene expression **[Table E3a],** enriched their association marginally with FVC (18.6% (n=725)), FEV_1_ (30% (N=1,172)), and TLC (16.2% (n=633)) but substantially with RV (0.15% (n=6)), FRC (4.3% (n=169)), and IC (2.5% (n=96)) **[Table E3b-g]**. These eQTLs were also enriched for their association with race/ethnicity among those associated with FVC (36.7% (n=266)), FEV_1_ (33.7% (n=395)), TLC (29.2% (n=185)) and IC (6.8% (n=60)), but not for FRC (5.3% (n=9)).

GO pathways for eQTLs associated with FVC, FEV_1_, TLC, and FRC included antigen presentation, lymphocyte differentiation, microtubule polymerization and bundle formation, and regulation of reactive oxygen species **[Fig. 1]**. eQTLs associated with FVC and FEV_1_ were additionally enriched for actin filament/ciliary dependent cellular secretory processes and movement, small GTPase-mediated signal transduction, muscle development, organic acid biosynthesis, while eQTLs associated with FRC and TLC were additionally enriched for fatty acid metabolism, lipid, specifically sphingolipid biosynthesis, chromosome segregation and mitosis, calcium ion homeostasis, and neural and epithelial tube formation **[Fig. 1]**. eQTLs associated with IC and RV did not encode for any pathway.

**Figure 1.**
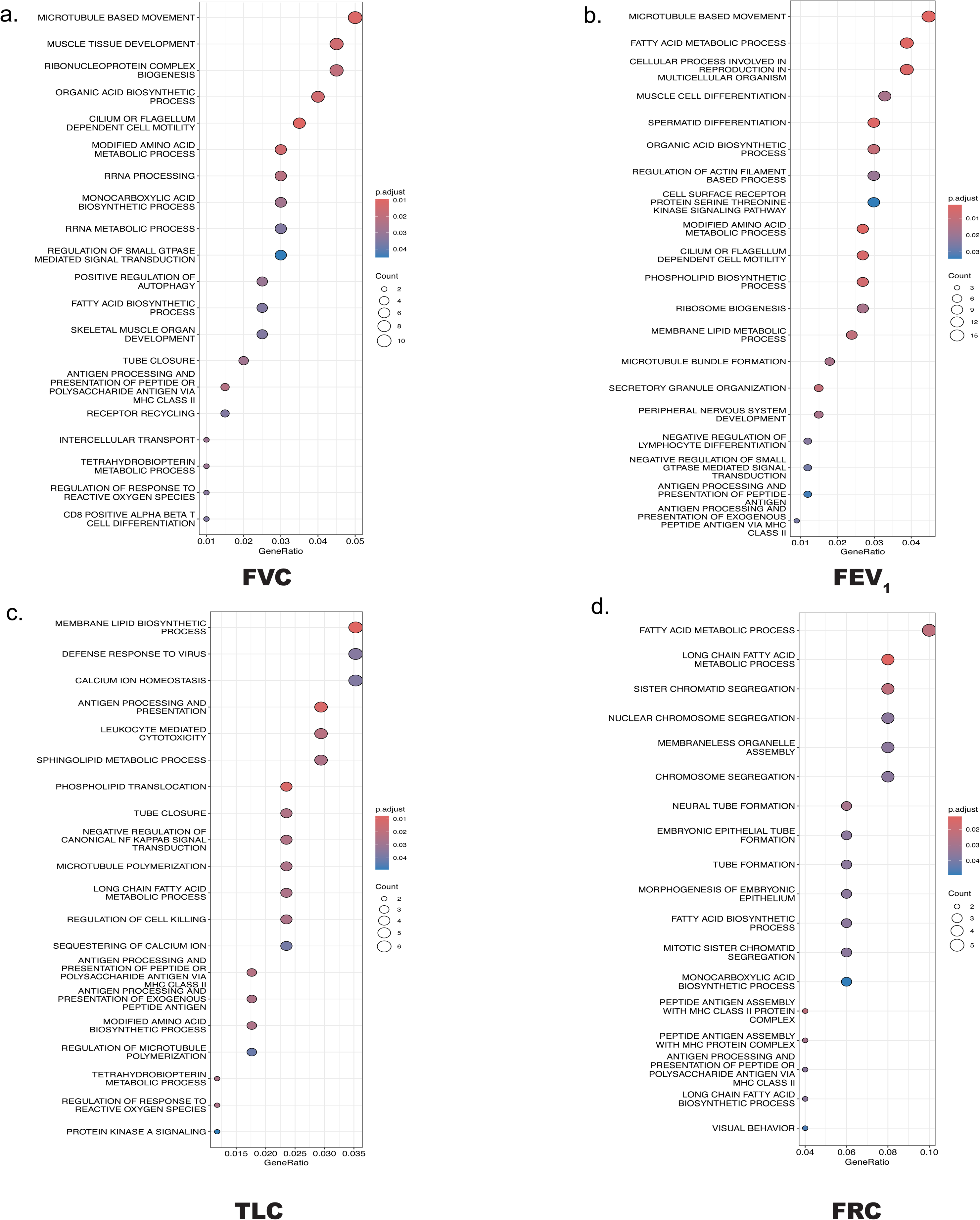
Enrichment of GO pathways in eQTLs associated with asthma burden. GO pathways for eQTLs associated with **a)** FVC, **b)** FEV_1_, **c)** TLC and **d)** FRC were enriched for antigen presentation, lymphocyte differentiation, microtubule polymerization and bundle formation, and regulation of reactive oxygen species. eQTLs associated with **a)** FVC and **b)** FEV_1_ were additionally enriched for actin filament/cilia dependent cellular secretory processes and movement, small GTPase mediated signal transduction, muscle development, organic acid biosynthesis, while eQTLs associated with **c)** FRC and **d)** TLC were additionally enriched for fatty acid metabolism, lipid, particularly sphingolipid biosynthesis, chromosome segregation and mitosis, calcium ion homeostasis, and neural and epithelial tube formation.

### Association of metabolic measures, anthropometrics, and Th cell subsets, with asthma burden and eQTLs

Among metabolic measures, triglyceride levels were associated with FVC, IC, and RV/TLC ratio, insulin and HOMA-IR were associated with FRC, ERV, and ACT, HDL was associated with ACT, leptin was associated with FEV_1_/FVC ratio, FRC, RV, and ERV, and adiponectin was associated with FRC **[Fig. 2a]**. While all anthropometric measures were associated with FEV_1_/FVC ratio, FRC, and IC, midarm, waist, and hip circumference, and BMI were associated with RV/TLC ratio **[Fig. 2b]**. Among Th cells, Th1 cells were associated with ERV, Th1/Th17 cells were associated with ERV and IC, Th17 cells were associated with FEV_1_, FEV_1_/FVC ratio, RV, TLC, FRC and ERV, and CASI score, and naïve/Th2 cells were associated with FEV_1_, FEV_1_/FVC ratio, RV, TLC, and ERV. Tregs were associated were FEV_1_/FVC ratio **[Fig. 2c]**. eQTLs were minimally associated with neck and midarm circumference **[Table E4a, b],** triglycerides, insulin and leptin **[Table E4c-e],** and substantially associated with Th cell subsets **Table E4f-i]**.

**Figure 2.**
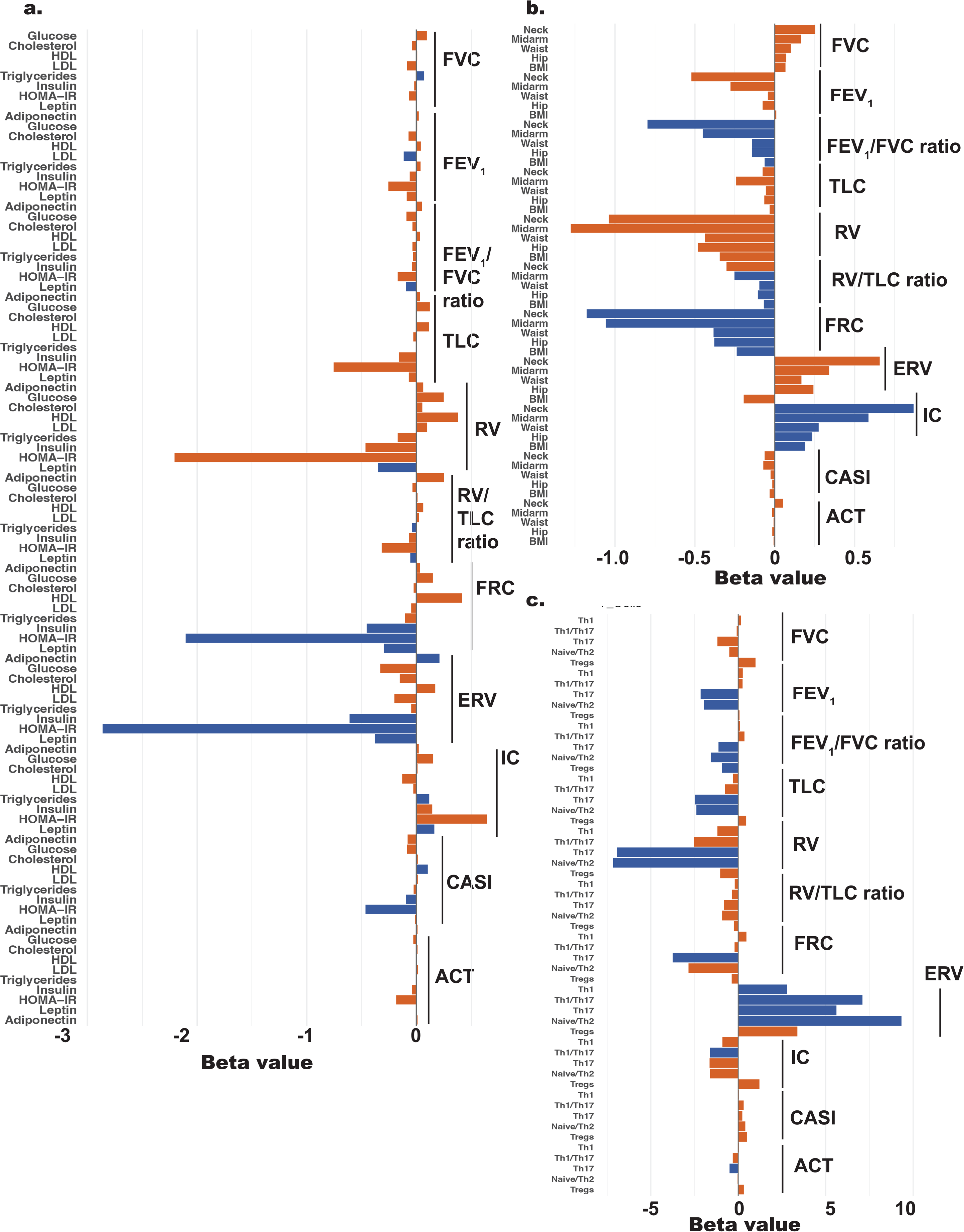
Association of anthropometric and metabolic measures, and Th cell proportions with asthma burden. The association of pulmonary function indices, CASI, and ACT with **a)** anthropometric and **b)** metabolic measures, and with **c)** Th cell proportions is summarized here. Significant associations are shown as blue bars, while associations that did not reach statistical significance are shown as orange bars.

### Attenuation of eQTL association with asthma burden by anthropometrics, metabolic measures, and Th cell subsets

Next, we investigated the effect of anthropometrics, metabolic measures, and Th cell subsets on the association of eQTLs with asthma burden. While 55.7% (405 of 725) eQTLs retained significant association with FVC, 44.2% were attenuated by waist circumference, leptin and adiponectin, and proportions of Th1, Th17 and Tregs **[Fig. 3a]**. Similarly, 42.4% (497 of 1172) eQTLs retained significant association with FEV_1_, while 47.6% were attenuated by neck, midarm, and waist circumference, leptin and adiponectin, and proportion of Th1, Th17, and Tregs **[Fig. 3b]**. The association of 46.7% (296 of 633) eQTLs with TLC retained significance, while 53.3% were attenuated by waist circumference, HOMA-IR and leptin, and Th1 and Th17 cell proportions **[Fig. 3c]**. The association of 50.3% (83 of 169) eQTLs with FRC retained significance, while 49.7% were attenuated by BMI, HOMA-IR, leptin and adiponectin, but not by any Th cell subset **[Fig. 3d]**. The association of 78.1% (75 of 96) eQTLs with IC retained significance, 21.9% were attenuated by waist circumference, and proportions of Th1 and Tregs **[Fig. 3e]**.

**Figure 3.**
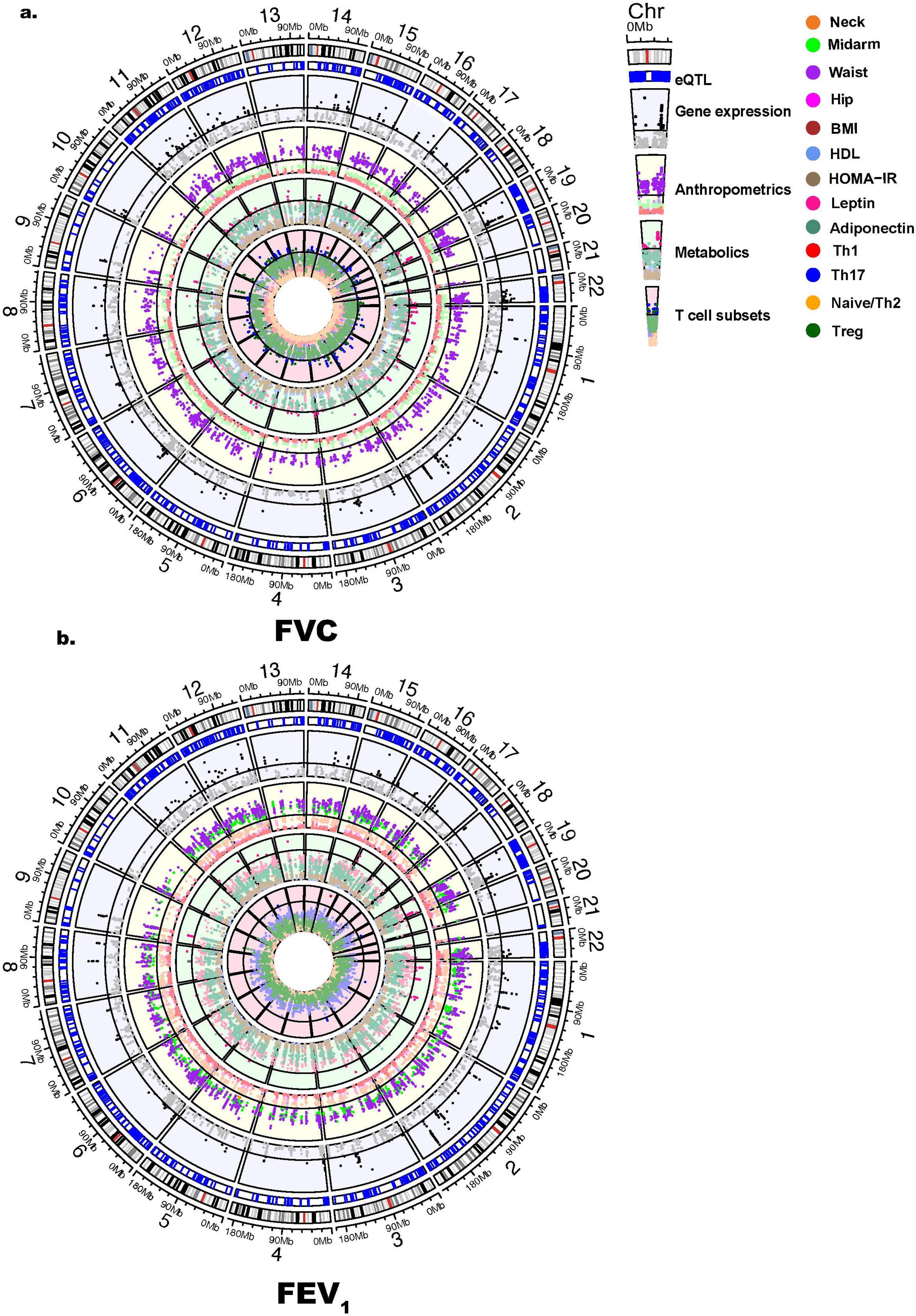
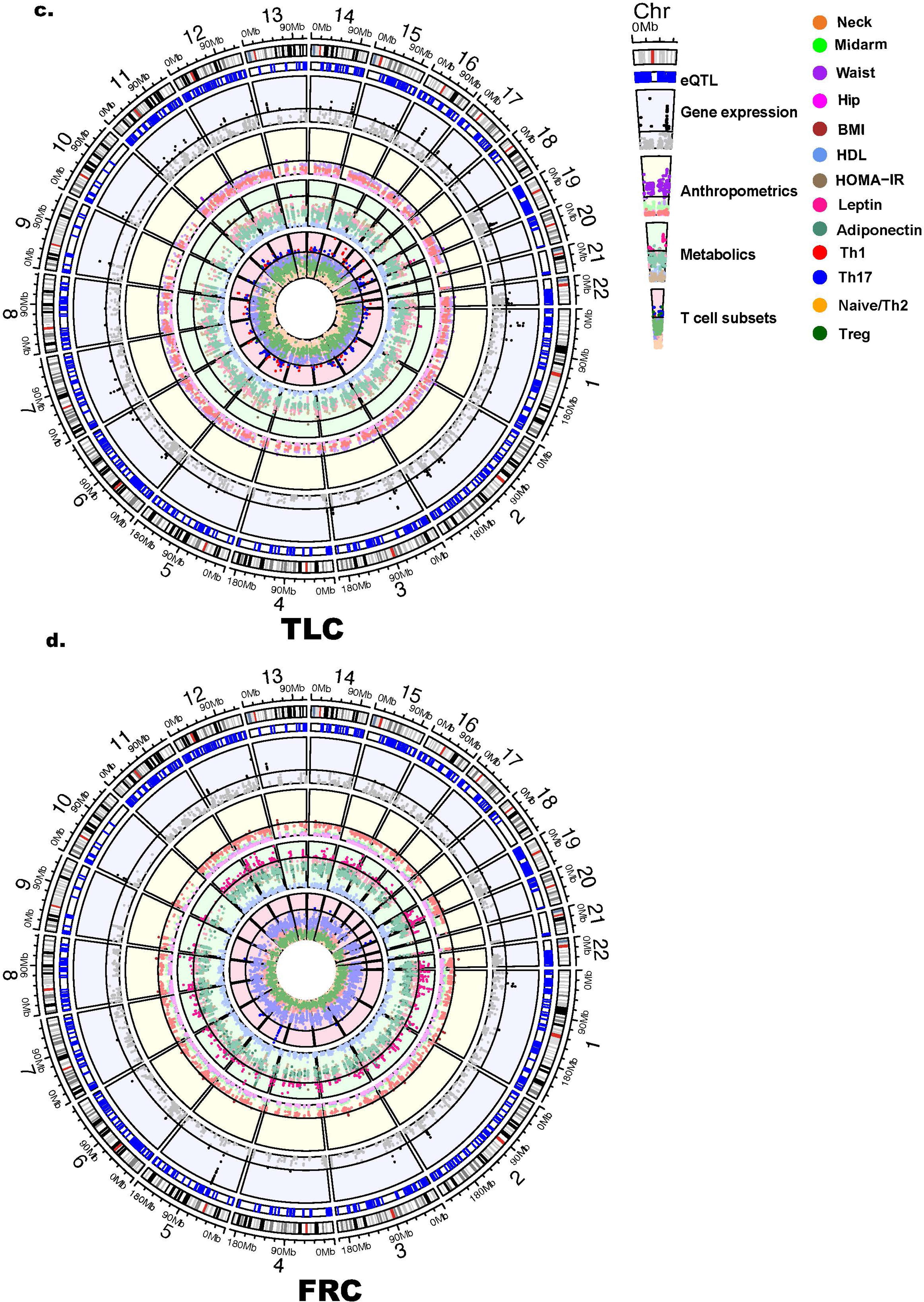
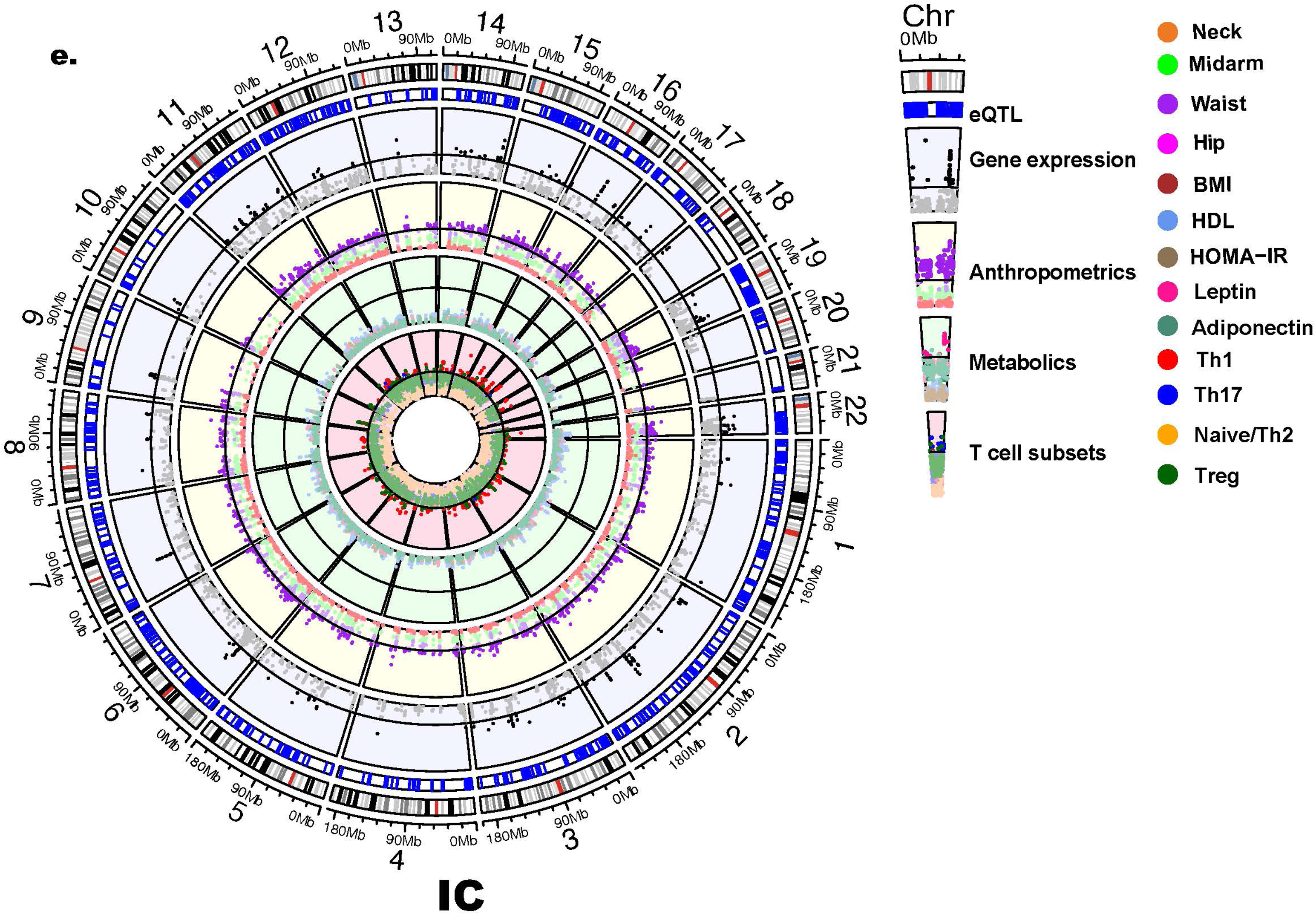
Association of eQTLs, expression of their corresponding genes, and of anthropometric and metabolic measures, and Th cell proportions with asthma burden. The circus plots comprehensively illustrate the variables associated with **a)** FVC, **b)** FEV1, **c)** TLC, **d)** FRC, and **e)** IC in multivariate analysis. Among all eQTLs shown as strokes in the outermost circle, the second circle illustrates the eQTLs (marked as blue strokes) that retained statistical significance in their association with each lung function index. For the remainder of the variables, expression of genes corresponding to the eQTLare in purple circles, anthropometric measures in yellow circles, metabolic measures in green circles, and Th cell proportions in pink circles. Among these, the outer circle in each set of colored circles reports significant associations while the inner circle report associations that do not reach statistical significance. For instance, in the outer purple circle, each black dot is a gene encoded by the corresponding eQTL that is significantly associated with the pulmonary function index, while in the inner purple circle, each grey dot is a gene encoded by the eQTL that is not significantly associated with the pulmonary function index. While the outer yellow circle denotes statistically significant association of waist (purple dot), hip (pink dot), midarm (green dot), and neck (orange dot) circumference with the pulmonary function indices, the outer green circle denotes statistically significant association of leptin (fuschia dot), adiponectin (forest green dot) and HOMA-IR (dark grey dot), and the outer pink circle denotes statistically significant association of Th17 cells (dark blue dot), Th1 cells (red dot), naYve/Th2 (yellow dot) and T regulatory cells (green dot) with pulmonary function indices independent of other variables, the inner circles of corresponding colors report on association of anthropometrics, metabolic measures and Th cell proportions that were not statistical significant.

The GO pathways attenuated in their association with FVC, FEV_1_, and TLC included organic acid biosynthesis, antigen presentation, lymphocyte differentiation, actin filament/ciliary dependent cellular secretory processes, and small GTPase mediated signal transduction. Pathways that retained significance included microtubule-based movement and response to reactive oxygen species. Among pathways associated with FRC, fatty acid metabolism and chromosome segregation and mitosis additionally retained significance **[Fig. 4a-d]**.

**Figure 4.**
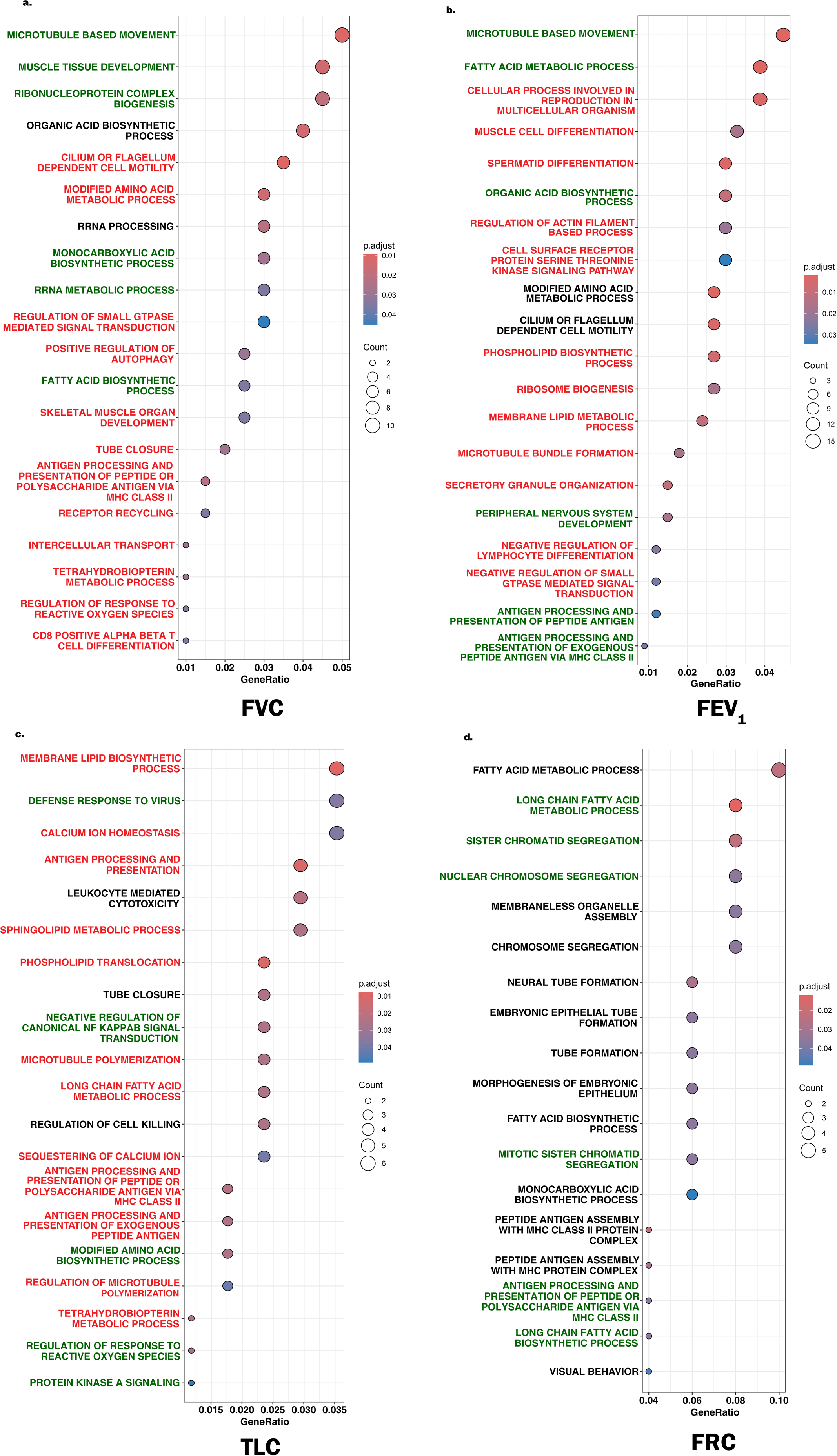
Effect of anthropometric and metabolic measures and Th cell proportions on GO pathways enriched in eQTLs associated with asthma burden. Among pathways enriched in **a)** FVC, **b)** FEV_1_ and **c)** TLC, those attenuated by anthropometric and metabolic measures and Th cell proportions are marked in red font – these included antigen presentation, lymphocyte differentiation, actin filament/ciliary dependent cellular secretory processes, and small GTPase mediated signal transduction; those that retained significance included fatty acid metabolism, organic acid biosynthesis and ribonucleoprotein complex biogenesis. Among pathways associated with **d)** FRC, no specific pathway was attenuated while those associated with fatty acid metabolism and chromosome segregation and mitosis retained significance.

Given the attenuating effects of anthropometric and metabolic measures and Th subset proportions on eQTL association with asthma burden, we applied SNF analysis, factoring in all four datasets to cluster participants with high disease burden, agnostic of their study group. The combination of datasets distinguished the highest number (5 of 10) of pulmonary function indices and clustered the study cohort into two clusters. Cluster 1, the cluster with lower pulmonary function was enriched for OA (66% (n=38)) relative to HwA (15% (n=5)), Ob (44% (n=12)) and HC (3.7% (n=3)). Compared to Cluster 2, Cluster 1 had worse anthropometric, metabolic and Th cell profiles in almost all groups, with highest differences in the OA group [**Table 2]**; the OA group had worse pulmonary function deficits as compared to other groups in Cluster 1 **[Fig. 5]**.

**Figure 5.**
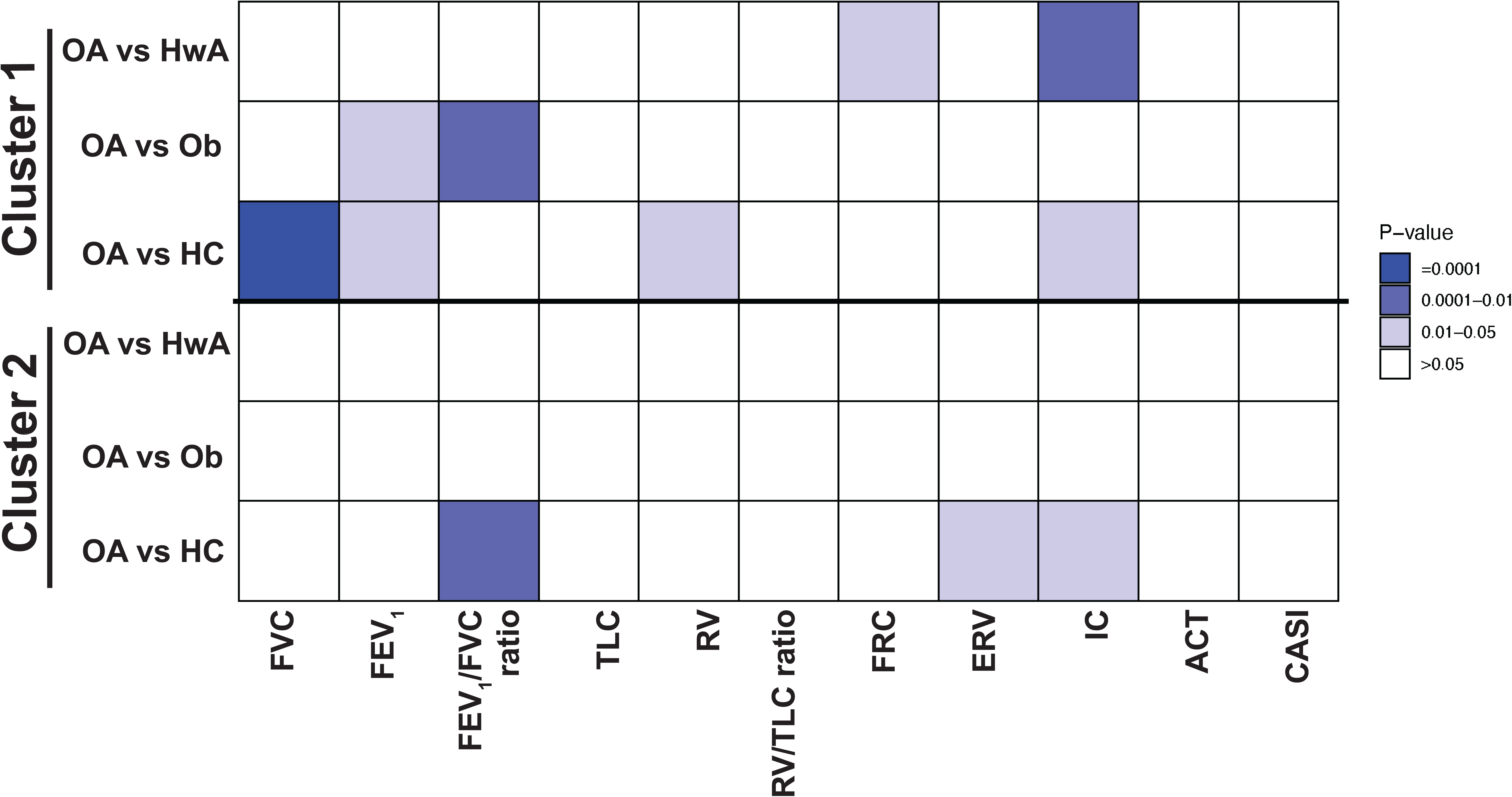
Comparison of lung function between the study groups in the two SNF clusters. Comparison of lung function indices between obese asthma group and the other three groups is summarized for samples in Cluster 1 and Cluster 2. There were more differences between obese asthma and healthy-weight asthma, obese no-asthma, and healthy-controls in Cluster 1 as compared to participants in each study group in Cluster 2.

**Table 2.** Comparison of pulmonary function and anthropometric, demographic, metabolic and immune measures between the clusters in each study group.

|  | **Obese asthma**<br>**(n=57)** |  | **Healthy-weight asthma**<br>**(n=33)** |  | **Obese no-asthma**<br>**(n=27)** |  | **Healthy weight controls**<br>**(n=27)** |  |
| --- | --- | --- | --- | --- | --- | --- | --- | --- |
|  | **Cluster 1**<br>**(n=38)** | **Cluster 2**<br>**(n=19)** | **Cluster 1**<br>**(n=5)** | **Cluster 2**<br>**(n=28)** | **Cluster 1**<br>**(n=12)** | **Cluster 2**<br>**(n=15)** | **Cluster 1**<br>**(n=3)** | **Cluster 2**<br>**(n=24)** |
| **Demographics** |  |  |  |  |  |  |  |  |
| Age (years) | 13.11 ± 2.40* | 8.84 ± 1.86* | 13.80 ± 1.30* | 10.11 ± 3.01* | 12.25 ± 1.49* | 10.20 ± 2.88* | 15.00 ± 2.65 | 10.96 ± 2.91 |
| Male (n (%)) | 13 (34) | 11 (58) | 4 (80) | 17 (61) | 4 (33) | 5 (33) | 2 (67) | 13 (54) |
| African<br>American<br>(n(%)) | 26 (68) | 12 (63) | 2 (40) | 19 (68) | 10 (83) | 9 (60) | 3 (100) | 11 (46) |
| **Pulmonary Function Indices \*\*** |  |  |  |  |  |  |  |  |
| FVC | 100.91 ± 17.73 | 101.26 ± 14.41 | 96.25 ± 9.91 | 98.42 ± 17.65 | 107.82 ±<br>14.56 | 99.50 ± 12.92 | 83.00 ± 1.73* | 96.83 ±<br>13.19* |
| FEV1 | 90.25 ± 20.40 | 96.89 ± 17.86 | 93.75 ± 13.55 | 95 ± 16.35 | 105.82 ±<br>10.72 | 98.14 ± 11.63 | 76.67 ± 11.85 | 96.96 ± 14.22 |
| FEV1/FVC | 78.91 ± 9.46* | 84 ± 6.43* | 85.25 ± 8.62 | 86.04 ± 8.50 | 87.55 ± 5.09 | 88.36 ± 5.09 | 84.33 ± 5.69 | 88.54 ± 6.01 |
| RV | 96.59 ± 33.67 | 119.67 ± 52.73 | 110.75 ±<br>37.72 | 132.61 ±<br>50.82 | 84.73 ± 37.23 | 132.91 ±<br>113.50 | 111.5 ± 3.54 | 105.77 ±<br>32.06 |
| TLC | 91.31 ± 16.26 | 99.89 ± 13.87 | 88.25 ± 4.99 | 97.42 ± 20.71 | 93.73 ± 17.27 | 101.27 ±<br>29.25 | 86 ± 1.41* | 95.32 ±<br>12.80* |
| RV/TLC | 23 ± 5.85 | 26.28 ± 8.94 | 26.75 ± 7.89 | 29.13 ± 8.69 | 19.18 ± 6.55 | 26.64 ± 12.40 | 27.5 ± 2.12 | 24.09 ± 5.86 |
| FRC | 79.75 ± 18.80 | 97.76 ± 23.58* | 94.5 ± 9.00* | 99 ± 25.54 | 76.64 ± 25.29 | 102.64 ±<br>55.50 | 94 ± 11.31 | 97.14 ± 17.68 |
| ERV | 69.53 ± 78.30 | 63 ± 21.14 | 62.25 ± 21.47 | 73.21 ± 42.49 | 63.55 ± 27.88 | 71.79 ± 27.91 | 101 ± 40.63 | 79.54 ± 24.75 |
| IC | 101.66 ± 25.68 | 101.11 ± 11.97 | 80.75 ± 7.23* | 93.68 ±<br>24.61* | 107.82 ±<br>13.45 | 100 ± 14.83 | 72 ± 11.14 | 91.5 ± 15.79 |
| CASI | 6.77 ± 3.72 | 8.57 ± 2.68 | 5.00 ± 4.00 | 7.35 ± 3.65 | NA | NA | NA | NA |
| ACT | 18.26 ± 5.42 | 17.37 ± 5.68 | 21.40 ± 4.56 | 19.39 ± 4.04 | 18.25 ± 7.41 | 19.38 ± 8.50 | 22 ± 3.00 | 18.36 ± 9.41 |
| **Anthropometric measures** |  |  |  |  |  |  |  |  |
| Neck | 36.54 ± 4.35* | 31.03 ± 3.40* | 33.80 ± 2.41* | 29.11 ± 2.60* | 35.04 ± 2.80* | 31.47 ± 2.57* | 34.00 ± 4.50 | 29.46 ± 2.37 |
| Midarm<br>(cms) | 35.06 ± 6.13* | 25.23 ± 3.02* | 26.90 ± 2.90* | 21.36 ± 3.12* | 34.81 ± 4.53* | 27.84 ± 3.52* | 25.00 ± 3.04 | 22.02 ± 3.01 |
| Waist (cms) | 103.28 ±<br>20.49* | 76.14 ± 11.24* | 76.50 ± 12.12 | 62.40 ± 7.44 | 96.75 ±<br>12.52* | 78.85 ±<br>10.58* | 68.83 ± 6.01 | 64.65 ± 8.31 |
| Hip (cms) | 112.62 ±<br>17.25* | 85.64 ± 9.07* | 94.88 ± 9.10* | 77.48 ±<br>10.20* | 109.46 ±<br>12.19* | 91.80 ±<br>11.16* | 90.67 ± 6.65 | 79.90 ± 10.70 |
| Height (cms) | 159.71 ± 9.91* | 136.01 ± 9.84* | 165.30 ±<br>10.14* | 141.61 ±<br>16.03* | 158.04 ±<br>9.17* | 145.16 ±<br>12.44* | 174.97 ±<br>17.84 | 148.80 ±<br>16.93 |
| Weight (kg) | 85.69 ± 25.23* | 44.14 ± 10.74* | 60.52 ±<br>12.30* | 36.51 ±<br>12.80* | 80.78 ±<br>18.60* | 52.62 ±<br>12.50* | 58.23 ± 10.16 | 41.45 ± 13.61 |
| BMI<br>percentile | 97.39 ± 3.54* | 95.57 ± 3.57* | 73.71 ±<br>25.63* | 51.27 ±<br>29.39* | 97.31 ± 3.46* | 94.69 ± 5.36* | 32.79 ± 21.31 | 52.79 ± 22.84 |
| **Metabolic measures** |  |  |  |  |  |  |  |  |
| Glucose<br>(mg/dl) | 80.34 ± 8.30 | 76.89 ± 6.57 | 89.20 ± 15.83 | 75.30 ± 8.26 | 82.64 ± 7.53 | 80.27 ± 7.62 | 79.67 ± 3.21 | 80.81 ± 6.49 |
| Cholesterol<br>(mg/dl) | 153.08 ± 29.66 | 157.58 ± 36.99 | 151 ± 33.96 | 148.59 ±<br>39.66 | 137.64 ±<br>23.94 | 138.27 ±<br>46.23 | 150 ± 18.36 | 151 ± 22.41 |
| HDL (mg/dl) | 45.13 ± 7.83* | 51.11 ± 11.38* | 46.78 ±<br>10.01* | 59.86<br>±13.53* | 38.55 ±<br>12.94* | 48.79 ± 9.24* | 41.73 ± 6.57* | 56.17 ±<br>11.89* |
| LDL (mg/dl) | 88.73 ± 24.24 | 86.79 ± 26.42 | 90.36 ± 33.48 | 76.87 ± 34.02 | 72.23 ± 24.02 | 70.89 ± 40.36 | 84.77 ± 14.70 | 99.45 ±<br>114.22 |
| Triglycerides<br>(mg/dl) | 93.82 ± 65.72 | 78.95 ± 41.83 | 58.40 ± 9.34 | 53.63 ± 16.95 | 109.82 ±<br>35.38* | 71.20 ±<br>25.90* | 62.33 ± 9.24 | 69.57 ± 27.59 |
| Insulin<br>(μU/ml) | 29.21 ± 22.27* | 11.26 ± 5.68* | 46.34 ± 70.51 | 8.04 ± 5.96 | 35.03 ± 22.29 | 18.13 ± 26.23 | 12.16 ± 13.55 | 8.26 ± 8.14 |
| HOMA-IR | 5.89 ± 4.70* | 2.17 ± 1.13* | 12.40 ± 20.02 | 1.55 ± 1.26 | 7.07 ± 4.30 | 3.86 ± 6.24 | 2.32 ± 2.51 | 1.62 ± 1.47 |
| Leptin<br>(ng/ml) | 60.01 ± 31.05* | 30.45 ± 24.35* | 16.46 ± 6.62 | 7.75 ± 11.19 | 51.87 ±<br>21.97* | 26.39 ±<br>17.29* | 5.33 ± 1.00 | 5.46 ± 6.27 |
| Adiponectin<br>(µg/ml) | 5.01 ± 2.56* | 7.50 ± 3.47* | 6.90 ± 3.27 | 8.99 ± 4.25 | 4.14 ± 1.79* | 8.24 ± 2.73* | 3.09 ± 2.88* | 8.92 ± 5.20* |
| **Percent T helper cell proportions** |  |  |  |  |  |  |  |  |
| Th1 | 7.88 ± 3.30 | 9.38 ± 3.63 | 10.14 ± 2.01 | 8.38 ± 4.38 | 8.63 ± 6.39 | 9.49 ± 4.57 | 12.63 ± 1.99* | 8.76 ± 3.37* |
| Th1/Th17 | 4.19 ± 2.48 | 3.68 ± 2.42 | 4.27 ± 1.11 | 4.08 ± 2.62 | 4.18 ± 3.25 | 4.28 ± 2.38 | 10.41 ± 2.71 | 4.20 ± 1.81 |
| Th17 | 4.41 ± 1.98* | 2.95 ± 0.94* | 5.24 ± 1.74* | 3.05 ± 1.08* | 4.30 ± 1.29* | 2.84 ± 1.16* | 5.03 ± 1.83 | 2.63 ± 1.19 |
| Naïve/ Th2 | 3.84 ± 1.64 | 3.04 ± 1.34 | 4.68 ± 0.77* | 2.97 ± 1.26* | 2.93 ± 0.94 | 2.23 ± 0.99 | 4.67 ± 0.58* | 2.63 ± 1.29* |
| Regulatory<br>T cells | 6.44 ± 1.78* | 7.86 ± 2.25* | 6.31 ± 1.51 | 6.52 ± 2.15 | 6.54 ± 1.38 | 5.47 ± 1.64 | 9.85 ± 1.28* | 6.14 ± 1.71* |

\*Between-cluster differences in each group that were statistically significant
All continuous variables are reported as mean ± SD, proportions are reported as n (%).
\*\* Percent predicted values of pulmonary function indices other than FEV<sub>1</sub>/FVC ratio and RV/TLC ratio are reported.

Moreover, MAF of 285 eQTLs differed between OA samples in the two clusters, of which 54% (n=155) were associated with differential gene expression **[Table E5a,b]**. Similar to 3,904 eQTLs, the 155 eQTLs were associated with FVC (15.5% (n=24)), FEV_1_ (54.2% (n=84)), TLC (24.5% (n=38)), and FRC (12.9% (n=20)). While anthropometric measures, particularly waist circumference attenuated the association of eQTLs with FVC **[Fig. 6a]**, leptin in addition to waist, neck and midarm circumference attenuated the association of eQTLs with FEV_1_ **[Fig. 6b]**. Lung volumes differed from FVC and FEV_1_, wherein leptin, and Th1 and Th17 cells attenuated the association of eQTLs with TLC **[Fig. 6c]**, while adiponectin along with leptin, and Th1 and Th17 cells attenuated the association of eQTLs with FRC **[Fig. 6d]**. Pathway analysis was not feasible on these few eQTLs. Specific genes including *HLA-DQ2, PLA2G4C*, and one locus on *FAM167A* were attenuated for FVC, FEV_1_, TLC, and FRC, while *FRMD4B* were additionally attenuated for FVC, FEV_1_ and FRC, *CCDC127* was attenuated for FVC and FEV_1_, and *AKREI2* were additionally attenuated for TLC. eQTL encoding for *MEI1* retained significance for FVC, FEV_1_, TLC, and FRC, and *ATF6* was significant for all but FRC. Additional eQTLs specific to FVC and FEV_1_ included *GSTCD, FUT10,* and *DNAAF11,* and those for TLC and FRC included *GJA3* and *AKR1C3*.

**Figure 6.**
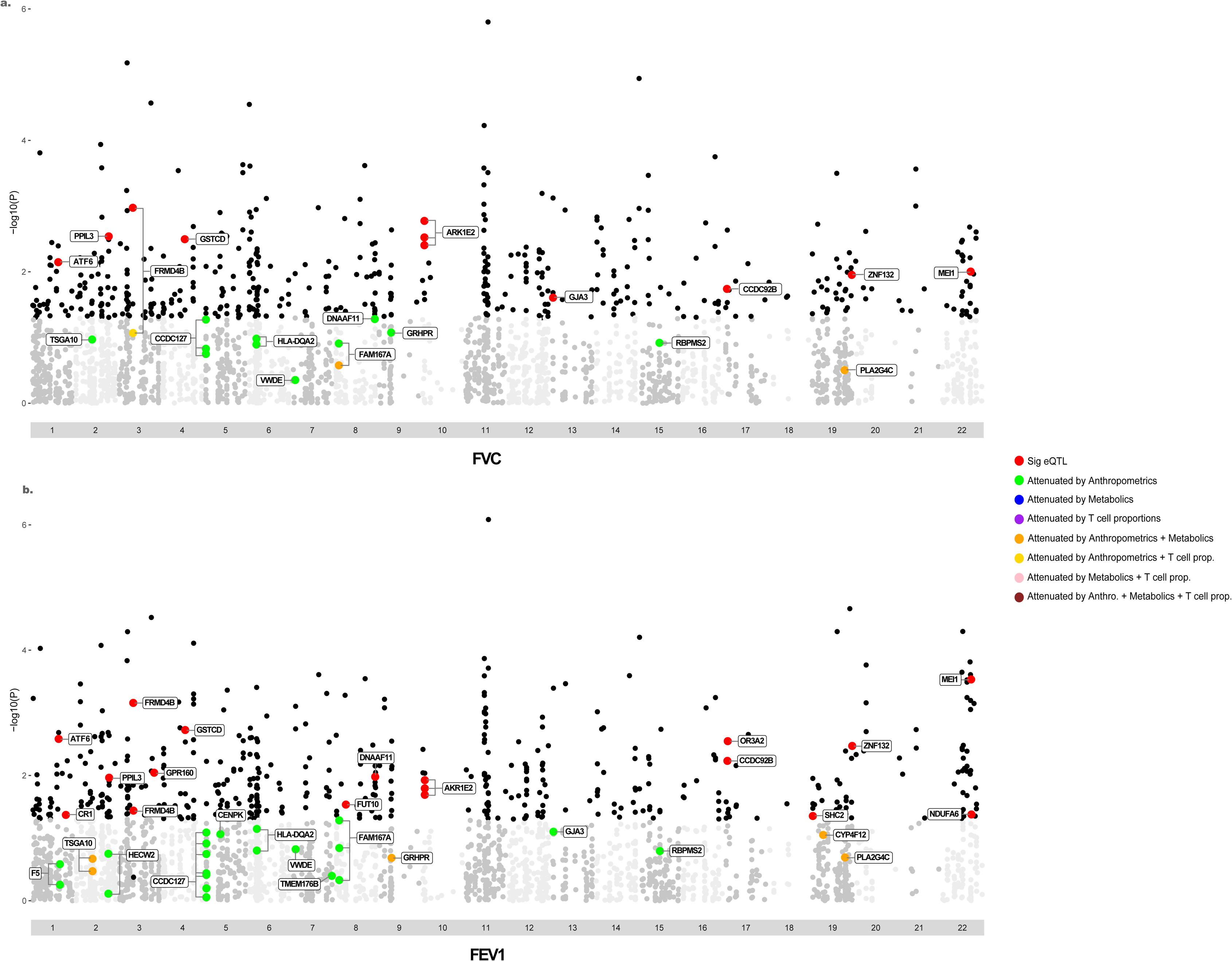
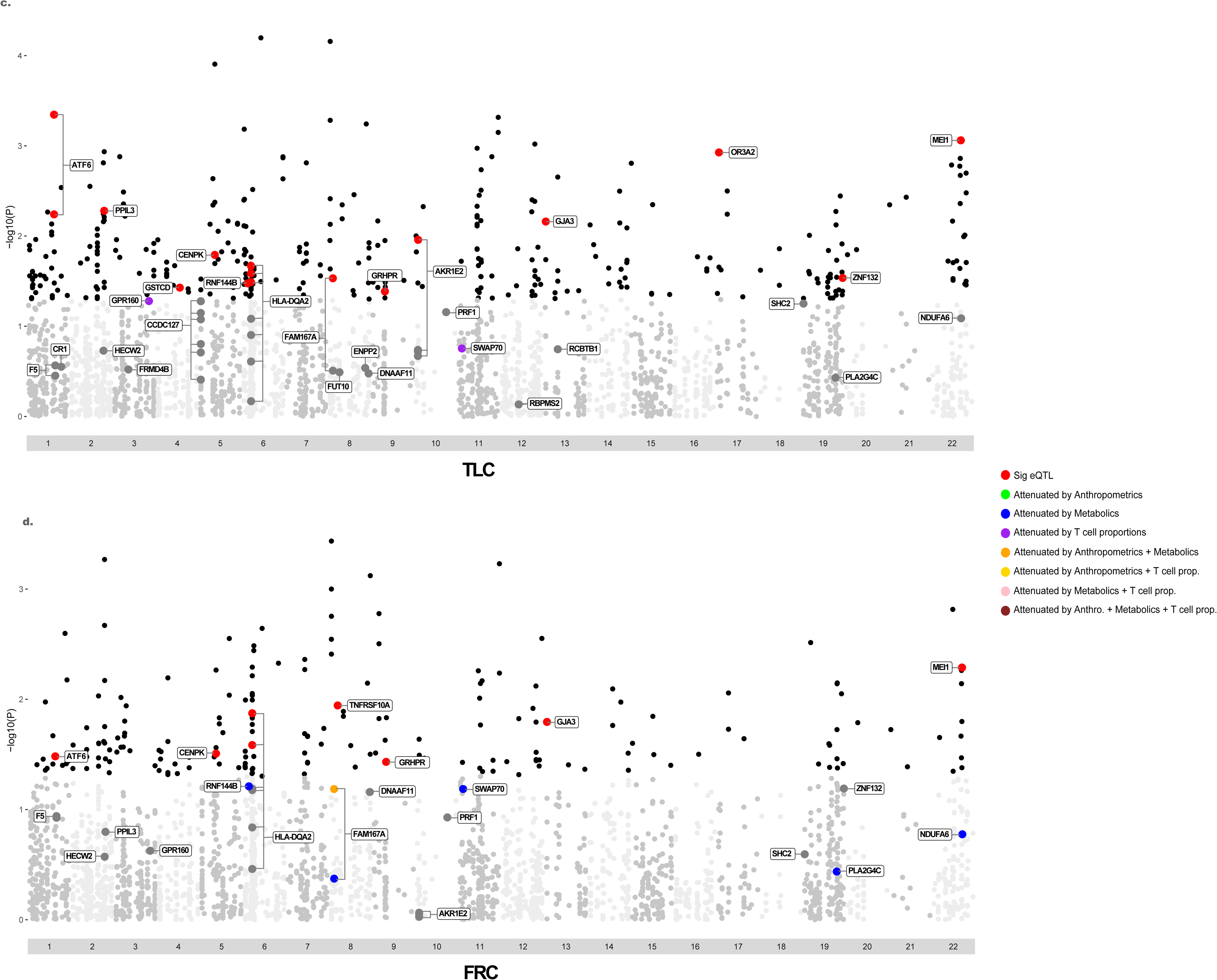
Attenuating effects of anthropometric and metabolic measures, and Th cell proportions on eQTLs associated with lung function indices in obesity-related asthma. The Manhattan plots illustrate the proportion of 155 eQTLs associated with **a)** FVC, **b)** FEV1, **c)** TLC, and **d)** FRC in obese asthma group that retained significance or were attenuated by anthropometric and metabolic measures and Th cell proportions. While red dots mark eQTLs that retained statistical significance for each pulmonary function index, green dots mark those attenuated by anthropometric measure(s) alone, blue dots mark those attenuated by metabolic measure(s) alone, purple dots mark those attenuated by Th cell proportion(s) alone, orange dots mark those attenuated by both anthropometric and metabolic measures, yellow dots mark those attenuated by anthropometric measure(s) and Th cell proportions, pink dots mark those attenuated by metabolic measure(s) and Th cell proportions, and maroon dots mark those attenuated by anthropometric and metabolic measures and Th cell proportions. eQTLs labeled in dark grey in plots on **c)** TLC and **d)** FRC are ones that were attenuated but did not retain significance with any anthropometric, metabolic measure or Th cell proportions.

### Ancestry of eQTLs associated with asthma burden

Given enrichment for racial differences in eQTLs associated with pulmonary function, between-cluster comparison revealed higher proportion of African ancestry in Cluster 1, the cluster with worse clinical profile **[Fig. 7]**. This enrichment was not driven by a specific study group.

**Figure 7.**
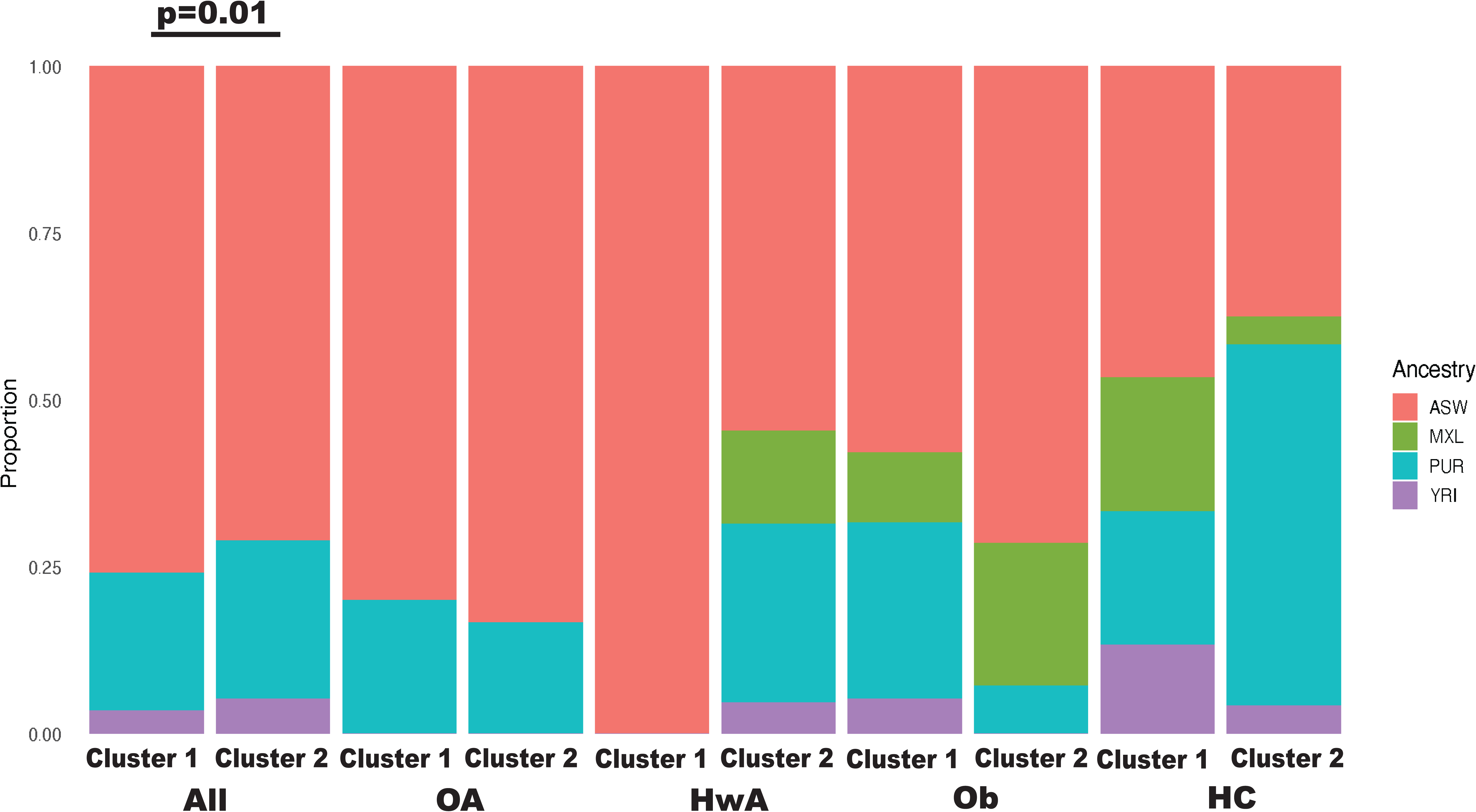
Ancestry of eQTLs associated with asthma burden. There were a higher proportion of children of African descent as compared to Mexican, Puerto Rican or Yorubian descent in Cluster 1, the cluster with higher disease burden. The higher proportion of African ancestry in Cluster 1 was not driven by any specific study group.

### Validation

We validated our observations in a second cohort comprised of 101 children ages 7-11 years **[Table 3]**. Of the 1,577 previously identified eQTLs in the validation cohort (18), 314 overlapped with 3,904 eQTLs in the primary cohort **[Table E6a]**. Of these, 12.1% (n=38) were nominally associated with FVC, 11.1% (n=35) with FEV_1_, 6.1% (n=19) with TLC, 4.8% (n=15) with FRC, and 7% (n=22) with IC **[Table E6b-f]**. Among these, while *RNASET2* was associated with FVC, FEV_1_, TLC and FRC*, FBLN5* was associated with FVC, FEV_1_, TLC and IC. An additional 26 genes overlapped between FVC and FEV_1_ among which *KIF16B, STX2, HEATR3,* and *SERPINB6* that have been associated with asthma and obesity.

**Table. 3.** Demographic and clinical characteristics of study participants in validation cohort.

|  | **Obese asthma**<br>**(n=48)** | **Healthy-weight asthma**<br>**(n=53)** | **P Value** |
| --- | --- | --- | --- |
| **Demographics** |  |  |  |
| Age (years) | 9.04 ± 1.34 | 8.85 ± 1.26 | 0.46 |
| Male (n (%)) | 27 (56) | 31 (58) | 0.98 |
| African American (n(%)) | 15 (32) | 15 (28) | 0.92 |
| **Pulmonary Function Indices\*** |  |  |  |
| FVC | 99.48 ± 14.49 | 92.26 ± 14.21 | 0.01 |
| FEV<sub>1</sub> | 91.69 ± 17.59 | 88.99 ± 15.82 | 0.42 |
| FEV<sub>1</sub>/FVC | 91.24 ± 9.58 | 95.81 ± 8.08 | 0.01 |
| RV | 97.21 ± 38.36 | 113.02 ± 30.01 | 0.03 |
| TLC | 94.85 ± 13.97 | 94.41 ± 11.85 | 0.87 |
| RV/TLC | 24.27 ± 7.25 | 29.00 ± 5.77 | <0.001 |
| FRC | 89.00 ± 22.60 | 105.22 ± 25.68 | 0.001 |
| ERV | 72.10 ± 19.29 | 91.53 ± 30.36 | <0.001 |
| IC | 98.54 ± 17.26 | 82.39 ± 13.43 | <0.001 |
| ACT | 19.87 ± 4.27 | 19.56 ± 4.60 | 0.72 |
| CASI | 8.81 ± 3.09 | 7.79 ± 3.45 | 0.12 |
| **Anthropometrics** |  |  |  |
| Neck (cms) | 30.89 ± 2.32 | 27.53 ± 2.14 | <0.001 |
| Midarm (cms) | 26.15 ± 3.78 | 20.05 ± 2.69 | <0.001 |
| Waist (cms) | 78.11 ± 10.63 | 61.28 ± 6.39 | <0.001 |
| Hip (cms) | 87.32 ± 13.14 | 73.97 ± 7.59 | <0.001 |
| Height (cms) | 139.90 ± 9.01 | 136.65 ± 9.60 | 0.08 |
| Weight (kg) | 47.28 ± 11.70 | 32.00 ± 7.19 | <0.001 |
| BMI percentile | 94.96 ± 4.70 | 55.28 ± 25.15 | <0.001 |

| **Metabolics** |  |  |  |
| --- | --- | --- | --- |
| Glucose (mg/dl) | 92.26 ± 8.42 | 88.18 ± 9.34 | 0.02 |
| Cholesterol (mg/dl) | 157.65 ± 30.80 | 154.89 ± 31.52 | 0.66 |
| HDL (mg/dl) | 50.98 ± 9.18 | 52.92 ± 14.08 | 0.41 |
| LDL (mg/dl) | 98.88 ± 29.60 | 91.84 ± 26.40 | 0.21 |
| Triglycerides (mg/dl) | 73.39 ± 33.27 | 69.54 ± 37.42 | 0.59 |
| Insulin ((μU/ml) | 17.01 ± 12.31 | 12.41 ± 6.81 | 0.02 |
| HOMA-IR | 3.96 ± 3.08 | 2.70 ± 1.54 | 0.01 |
| Leptin (ng/ml) | 19.98 ± 11.17 | 7.55 ± 5.81 | <0.001 |
| Adiponectin (μg/ml) | 14.60 ± 6.06 | 18.22 ± 8.12 | 0.01 |

All continuous variables are reported as mean ± SD, categorical variables are reported as proportions (n (%))
\* Percent predicted values of pulmonary function indices other than FEV<sub>1</sub>/FVC ratio and RV/TLC ratio are reported.

## Discussion

We report on genetic susceptibility for pediatric obesity-related asthma, as measured by the association of eQTLs, or functionally relevant SNPs, with asthma burden, and the modifying influences of obesity-mediated complications that are biological sequelae of obesogenic environment, on this relationship. eQTLs were associated with FVC, FEV_1_, TLC, FRC, RV, and IC but not with ERV, FEV_1_/FVC ratio and RV/TLC ratio, or symptom-based classification of asthma severity and control. In keeping with prior reports (14), obesity-mediated complications including adiposity, and metabolic and immune perturbations, were also associated with asthma burden. Among these, truncal adiposity, insulin resistance, leptin and adiponectin levels, and Th1 and Th17 cell proportions, attenuated the association of approximately half of the eQTLs with pulmonary function suggesting that genetic susceptibility for asthma burden is influenced by biological sequelae of obesogenic environment. This observation was verified by a study-group agnostic analysis that revealed enrichment of participants with obesity-related asthma among those with lower pulmonary function. We verified our findings in a validation cohort where several eQTLs found in the primary cohort were associated with lung function indices in the validation cohort, albeit nominally. Overall, eQTLs associated with asthma burden were enriched for African ancestry. Together, we report on novel genetic markers of pediatric obesity-related asthma that are enriched for African Ancestry and are influenced by obesity-mediated complications.

There was substantial overlap in biological processes encoded by eQTLs associated with spirometric indices, FEV_1_ and FVC, and with lung volume indices, TLC and FRC. These can be grouped into pathways encoding for T cell functions such as antigen presentation, lymphocyte differentiation, and microtubule polymerization, linked with cell mobility or chemotaxis, and pathways encoding for ubiquitous cellular functioning including organic acid metabolism, chromosome segregation, and ribosomal RNA processing. Since eQTLs were quantified for CD4+T cell transcriptome, their enrichment for T cell relevant pathways verifies cell-specific effects of eQTLs (33) defining genetic underpinnings for the role of T cells in pediatric obesity-related asthma (13, 34). Enrichment of eQTLs in microtubule polymerization and GTPase signal transduction process pathways is particularly relevant in the context of our prior reports on the role of CDC42, a RhoGTPase, in the pathobiology of obesity-related asthma (18, 21). Having found uninhibited chemotaxis of CDC42-enriched CD4+T cells and their crosstalk with airway smooth muscle (ASM) (35), our findings identify a role of genetic susceptibility in Th cell-ASM crosstalk in obesity-related asthma.

Exemplifying gene by environment interaction, pathways attenuated by obesity-mediated complications were specific to T cell function. While some eQTLs were attenuated by truncal adiposity alone, others were attenuated by truncal adiposity along with metabolic abnormalities and non-atopic Th1 and Th17-mediated inflammation. The latter observation, in the setting of higher proportion of naïve/Th2 cells in children with obesity-related asthma, suggests that mechanistic differences, such as enrichment of T cell functions, rather than cell proportions alone, have higher relevance in obesity-related asthma. Furthermore, the attenuating effects of metabolic measures, insulin resistance and leptin, that have been epidemiologically linked with obesity-related asthma (6) identifies a novel mechanism by which metabolic measures may influence disease burden in obesity-related asthma. Together, we highlight the complex interaction of genetic susceptibility with biological sequelae of obesogenic environments in obesity-related asthma. Once verified, our observations support targeted investigations of SNPs encoding for most differentially expressed genes that were modulated by truncal adiposity, metabolic or immune abnormalities as genetic at-risk markers for obese children at risk for asthma.

Specific genes, eQTLs encoding for *HLA-DQ2, PLA2G4C*, and one locus on *FAM167A* were consistently attenuated, each of which has a role in cellular functioning wherein their inhibition is associated with pro-inflammatory responses. *HLA-DQ2* plays a role in antigen presentation and CD4+ T cell tolerance, and *PLA2G4C* encodes for phospholipase A2-γ enzyme that metabolizes prostaglandins into leukotrienes with established roles in asthma (36), *FAM167A* increases genetic predisposition for autoimmune diseases, including rheumatoid arthritis and Sjogren syndrome, and its encoded protein associated with autoimmunity is highly expressed in bronchial epithelium (37). In contrast, eQTLs encoding for *ATF6* and *MEI1* were not influenced by obesity-mediated complications identifying two genes that confer independent genetic risk for obesity-related asthma. *ATF6*, encoding for Activation Transcription Factor 6, activates genes in the unfolded protein response (UPR) that is key to endoplasmic reticulum stress response and pro-inflammatory T cell responses (38). ATF6 activation in Th2 and Th17 cells is associated a mixed granulocytic airway inflammation that is reported in adult obesity-related asthma (39). *MEI1*, encodes for Miz1, a transcription factor, that when inhibited, induced Th1 response with augmented IL-12 expression in a murine model of asthma (40).

Among eQTLs that overlapped between primary and validation cohort, those encoding for six genes, *RNASET2, FBLN5, KIF16B, STX2, HEATR3,* and *SERPINB6* were associated, although nominally, with more than one lung function index in the validation cohort. While *RNASET2* encodes for T2 ribonuclease which is associated with T2 inflammation (41) and with lipotoxicity in obesity (42), *FBLN5*, encodes for fibulin-5 which upregulates integrins and cell junction proteins, including yes-associated protein (YAP) (43). Although fibulin-5 in CD4+T cells is not mechanistically linked with asthma or obesity, YAP and CDC42 influence each other (44). We therefore speculate that fibulin-5 may play a role in YAP interaction with CDC42 in CD4+ T cell crosstalk with ASM. *KIF16B* encodes for Kinesin Family Member 16B which facilitates microtubule-dependent intracellular transport and is associated with BMI in GWAS (45) but its association with asthma is not known. In keeping with the overlap between T1 and T2 responses, *STX2* encodes for syntaxin-2 which is associated with eosinophilic degranulation and severe asthma (46) as well as dysregulation of insulin release (47). The two other genes, *HEATR3*, a part of NOD2 signaling pathways, and *SERPINB6* encoding for intracellular protease inhibitors, are novel findings from our study and have not been previously linked to CD4+ T cells or obesity or asthma.

We further found that eQTLs associated with lower pulmonary function were enriched for African ancestry, verifying observations from Genes-Environments and Admixture in Latino Americans (GALAII) and Study of African Americans, Asthma, Genes and Environment (SAGEII) studies (48). However, there was no overlap with a curated list of genes associated with lung function (49), suggesting the eQTLs in our study are likely unique to obesity-related asthma, and distinct from all-comer asthma included in prior studies.

While we identify several novel eQTLs associated with obese asthma disease burden, including those influenced by obesity-mediated complications, there are limitations to our study. Our cohort was of a modest size and enriched for children with obesity-related asthma, which may skew our observations towards obesity and its complications. We addressed this concern by using SNF analysis which was agnostic to study groups. Our sample was also enriched for individuals who classified themselves as African American. While it is not reflective of the general population, given how few GWAS have been enriched for African Americans, our reports on SNPs specific to the African ancestry associated with asthma likely explains the lack of overlap between previously reported SNPs and of high relevance for a race that is studied to a limited extent. Moreover, the eQTLs in the validation cohort only reached nominal significance in their association with pulmonary function, which we speculate may be explained by the younger age of participants in the validation cohort.

In summary, we report novel markers of genetic susceptibility for obesity-related asthma, and attenuation of approximately half of them by obesity-mediated complications that are biological sequelae of obesogenic environment. These findings are direct evidence of gene by environment interaction suggesting that genetic susceptibility to asthma is amenable to modification by preventing obesity-mediated complications such as truncal adiposity, and metabolic and systemic immune perturbations. Moreover, the influence of truncal adiposity alone for some eQTLs and that of truncal adiposity along with leptin and insulin resistance and Th cell proportions on other eQTLs reinforces the complex interaction between myriad complications of obesity that together contribute to the obese asthma phenotype. Genes that retain independent association with pulmonary function, once verified in future studies, will serve as at-risk markers for asthma among children with obesity.

## Author contributions

DAT conducted all computational analysis and contributed to the conception of the manuscript, YBW and SD contributed to data acquisition, AR was responsible for cell-based experiments and analysis of T cell proportions. DR conceptualized and drafted the manuscript. DAT, YBW, SD, and AR reviewed, edited and approved the manuscript. The authors have declared no conflict of interest.

---

**Figure E1.**
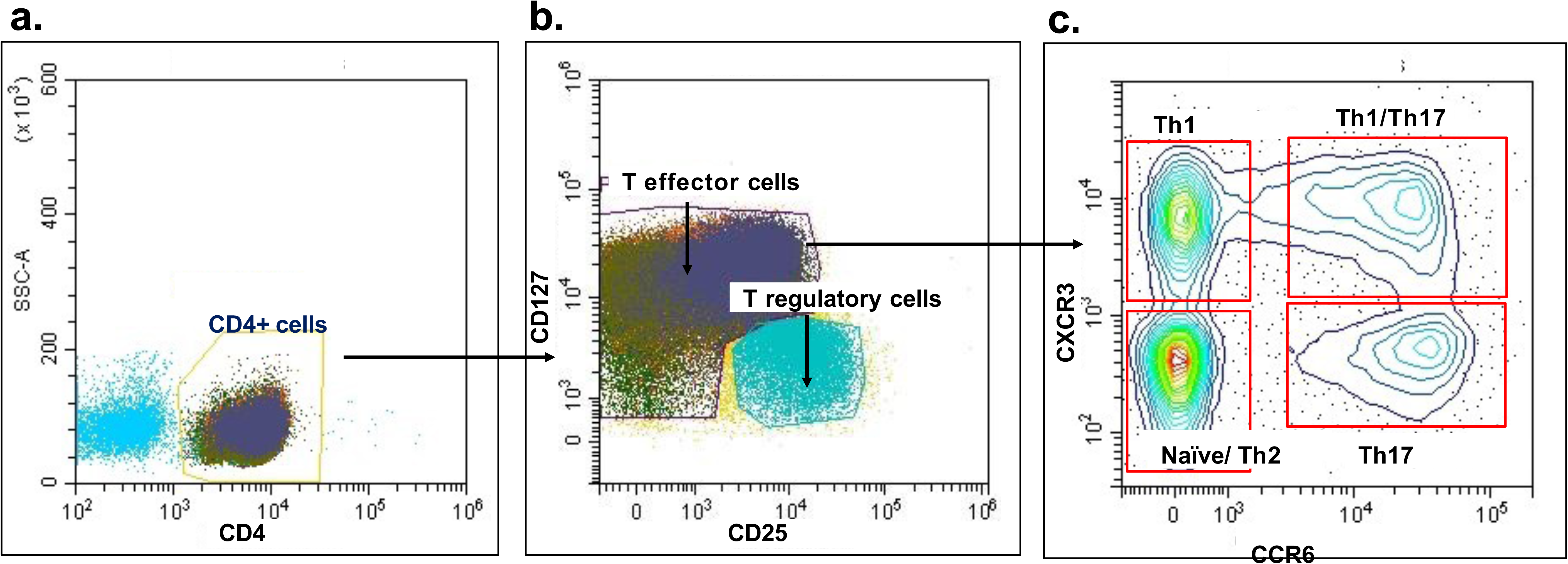
Schema of flow cytometric analysis of proportions of Th cell subsets. After gating on singlets, **a)** cells that stained for CD4 were identified, from which **b)** CD4+ cells were classified as regulatory and effector T cells based on CD25 and CD127 staining. **c)** Using CXCR3 and CCR6 staining, the effector T cells were further classified as Th1 cells (CXCR3+CCR6-), Th1/Th17 cells (CXCR3+CCR6+), naïve/Th2 cells (CXCR3-CCR6-), and Th17 cells (CXCR3-CCR6+).

