## Supplementary material for "Genetic susceptibility to obesity-related asthma and its modulation by sequelae of obesity": Online supplement

**Methods**

Study population

The primary study cohort, recruited between June 2019 and June 2023, was comprised of 144 children, ages 7-18 years. This cohort included 57 children with obesity-related asthma (OA), 33 with healthy-weight-asthma (HwA), 27 with obesity-alone (Ob), and 27 that were healthy-weight without asthma (healthy-weight controls (HC)). The validation cohort, recruited between July 2013 and August 2016 comprised of 101 children, ages 7-11 years, including 48 with obesity-related asthma (OA) and 53 with healthy-weight-asthma (HwA) (1). Obesity was defined as body mass index (BMI) >95^th^ percentile for age and sex (2). Asthma was defined by physician diagnosis in electronic medical records, active prescription of asthma medications, and reversible airflow obstruction (3). Informed consent was obtained from all participants and their primary caregivers on the date of the study visit, when they underwent anthropometric measurements, pulmonary function testing, and a fasting blood draw (4). The study was approved by the Institutional Review Boards at Albert Einstein College of Medicine and Children’s National Hospital, the two sites of study recruitment.

Measures of asthma disease burden

Asthma burden was measured using pulmonary function indices and symptom-based classification of severity using Composite Asthma Severity Index (CASI) (5) dichotomized to distinguish mild from moderate to severe asthma (CASI≤3), and control using Asthma Control Test (ACT) dichotomized to distinguish presence from absence of control (ACT<19) (6, 7). Of the pulmonary function indices, percent predicted values of forced vital capacity (FVC), forced expiratory volume in 1^st^ second (FEV_1_), and FEV_1_/FVC ratio were calculated using race-neutral Global Lung function Initiative (GLI)(8), and percent predicted values of mid expiratory flow rates (FEF_25-75%_), total lung capacity (TLC), residual volume (RV), RV/ TLC ratio, expiratory reserve volume (ERV), functional residual capacity (FRC) and inspiratory capacity (IC) were calculated using ATS/ERS guidelines (9).

Demographic, anthropometric, and metabolic variables

Demographic variables included age, sex, race/ethnicity, and anthropometrics included BMI percentile, and neck, midarm, waist, and hip circumference. Metabolic measures including glucose, insulin, total cholesterol, triglycerides, high density (HDL) and low-density (LDL) lipids, and adipokines, leptin and adiponectin, were quantified in fasting serum as previously described (10). Insulin resistance was defined by Homeostatic Measurement of Insulin Resistance (HOMA-IR) calculated as (glucose (mg/dl) * insulin ((μU/ml))/405.

T helper cell subsets

T helper cells were isolated from peripheral blood mononuclear cells separated from blood with Ficoll Hypaque density centrifugation method (4). Flow cytometric quantification of cell surface staining with CD4, CD25, CD127, CXCR3, and CCR6 antibodies (Biolegend, San Diego, US) on Beckman Coulter CytoFLEX Flow cytometer was analyzed on Flo Jo v.10.10.0 to quantify proportions of T helper cell subsets (Th1, Th1/17, naïve/Th2, Th17, and regulatory T cells) **[Figure S1]**.

Transcriptomic analysis of CD4+ T cells

500 ng RNA from CD4+T cells underwent library preparation with KAPA stranded mRNA-Seq kit (KAPA Biosystems, Wilmington, MA) that were sequenced on NovaSeq 6000. After quality analysis, transcripts were aligned to hg38 assembly using STAR aligner (v2.4.2a)(11), annotated using GenCode (v19), and analyzed on DESeq2. Normalized gene counts were analyzed using linear regression modeling to elucidate between-group differences, adjusting for covariates associated with gene expression variance (4).

Genotyping and eQTL mapping

CD4+ T cell DNA was genotyped on the Infinium Multi-Ethnic Global Bead Chip enriched for genetic variants in ethnically diverse populations (12). As previously reported (1), genetic variants were converted to the Watson (+) strand using the Genome build and Allele definition Conversion Tool to correspond to the transcriptome. Cis eQTLs were identified as variants within 1 Mb of a gene’s transcription start site that were associated with its expression. Analyses were limited to autosomal variants, removing those with a minor allele frequency (MAF) of less than 0.1 or those that did not follow the Hardy-Weinberg equilibrium (p-value<0.000001). Adjusted p-values were calculated by running 10,000 permutations using QTLtools cis, and the final eQTL set was defined by a conditional pass. To identify eQTLs in linkage disequilibrium (LD), the SNP2GENE module from Functional Mapping and Annotation of Genome-Wide Association Studies (FUMA GWAS) (13) was run on the final eQTL set to identify the lead SNPs and potential SNPs in LD with lead SNPs. The least stringent p value cut off for the module (1x10^-5^) was used to identify the maximum potential lead SNPs. Bulk RNA-seq data are available at GEO repository GSE227915 and genotype data on dbGap (database of Genotypes and Phenotypes) study identifier phs004474.v1.p1.

Statistical analysis

Analysis was conducted on R statistical software v. 4.4.1. First, we quantified the univariate association of eQTLs with pulmonary function indices, CASI, and ACT, limiting to those associated with expression of their corresponding gene. Gene ontology (GO) pathway analysis (14) was applied to elucidate the biological relevance of eQTLs associated with asthma burden. We applied Student T tests or Pearson correlation to quantify univariate association of anthropometrics, metabolic measures, and Th cell proportions with asthma burden. Using linear regression analysis, we quantified the effect of anthropometrics, metabolic measures, and Th cell proportions on association of eQTLs with asthma burden. Given association of eQTLs and anthropometrics, metabolic measures, and Th cell proportions with asthma burden, we applied Similarity Network Fusion (SNF) multi-omics analysis (15) to cluster samples with high disease burden irrespective of study group, since SNF assigns equal weightage to all datasets irrespective of their dimensionality. We also elucidated the ancestry of eQTLs associated with asthma burden with ADMIXTURE v1.3.0 (16). Finally, we investigated the overlap between eQTLs in primary cohort and the validation cohort, and the association of these overlapping eQTLs with pulmonary function, CASI and ACT in the validation cohort. False discovery rate (FDR) for all analyses was calculated using the Benjamini Hochberg method (17).
